# In-vivo subdermal implant in the alpha-Gal knockout mice model for the assessment of bioprosthetic heart valve calcification rate before and after a polyphenols-based treatment

**DOI:** 10.1101/2023.02.21.529461

**Authors:** Filippo Naso, Andrea Colli, Peter Zilla, Antonio Maria Calafiore, Chaim Lotan, Massimo A Padalino, Giulio Sturaro, Alessandro Gandaglia, Michele Spina

## Abstract

**Rationale:** The presence of preformed antibodies against αGal in the human lead to opsonization of the implanted bioprosthetic heart valve (BHV), leading to deterioration and calcification. Murine subcutaneous implantation of BHVs leaflets has been widely used for testing the efficacy of anti-calcification treatments, however, unlike the situation in humans, leaflets implanted into a murine model will not be able to elicit an αGal immune response because both donor and recipient species constitutively express the epitope.

**Objective:** This study evaluates the calcium deposition on commercial BHV using a new humanized murine αGal knockout (KO) animal model. Furtherly, the anti-calcification efficacy of a polyphenol-based treatment was deeply investigated.

**Methods and Results:** By using CRISPR/Cas9 approach an αGal KO mouse was created and used for the evaluation of the calcific propensity of original and polyphenols-treated BHV by subcutaneous implantation. The calcium quantification was carried out by plasma analysis; the immune response evaluation was performed by histology and immunological assays. Anti-αGal antibodies level in KO mice increases at least double after 2 months of implantation of original commercial BHV compared to WT mice, conversely, the polyphenols-based treatment seems to effectively mask the antigen to the KO mice’s immune system. Commercial leaflets explanted after 1 month from KO mice showed a four-time increased calcium deposition than what was observed on that explanted from WT. Polyphenol treatment prevents calcium deposition by over 99% in both KO and WT animals.

**Conclusions:** The implantation of commercial BHV leaflets significantly stimulates the KO mouse immune system resulting in massive production of anti-Gal antibodies and the exacerbation of the αGal-related calcific effect if compared with the WT mouse. The polyphenol-based treatment applied in this investigation showed an unexpected ability to inhibit the recognition of BHV xenoantigens by circulating antibodies completely preventing calcific depositions compared to the untreated counterpart.

## INTRODUCTION

Aortic valve disease is one of the most common valvular pathologies [1] with a significantly high mortality rate in symptomatic patients [2]. The most common cause of aortic valve disease in elderly patients (60-80 years old) is calcific degeneration [3]. The treatment of choice is surgical (SAVR, surgical aortic valve replacement) or transcatheter replacement (TAVR, transcatheter aortic valve replacement) according to guidelines [4]. Bioprosthetic heart valves (BHV, also known as “tissue valves”) are the most used type of device in more than 80% of all cases worldwide [3]. BHVs are made of bovine or porcine pericardium or leaflets valves and are conventionally cross-linked with glutaraldehyde (GA) to ensure tissue stability, reduce antigenicity, and maintain tissue sterility. They are mainly used in patients older than 60 years of age where the durability of the valve exceeds the life expectancy [5]. The functioning of BHVs is limited by shorter durability in younger patients in whom the process of calcification is accelerated [6] representing one of the limiting factors for their clinical application.

Traditionally, BHVs calcification has been attributed to extrinsic factors such as the chemical instability of GA, mechanical failure, and intrinsic ones like collagen degradation and calcium precipitation by residual lipids [7,8]. In recent years, the immune-mediate intrinsic pathway gained importance not least because of a better understanding of the αGal xenoantigen trigger [9–11]. The residual presence of αGal xenoantigen increases the human anti-galactose titers, starting from day 10 following BHV implantation [12] while reaching a peak at around 3 months [13] for IgM and IgG isotype. This sugar moiety is expressed in most mammalian tissues, except humans and higher primates. In humans, the continuous antigenic stimulation by gastrointestinal flora (expressing the αGal epitope) results in the production of anti-αGal antibodies accounting for 1 to 3% of the circulating immunoglobulins. Different research groups [10,14] have demonstrated that these preformed antibodies could cause opsonization of the valve tissue with consequent initiation of specific Fc-receptor-mediated macrophage recruitment with antigen processing and presentation, resulting in extracellular matrix (ECM) calcification and deterioration.

Murine subcutaneous implantation of BHVs leaflets has been widely used as an initial step for testing the efficacy of anti-calcification treatments [15–18]. However, unlike the situation in humans, bovine pericardial leaflets tissues implanted into a murine model will not be able to elicit an anti-Gal immune response because both donor and recipient species constitutively express αGal epitopes. Some studies have tried to demonstrate the link between the presence of the αGal antigen and the propensity to tissue calcification by comparing pericardial tissues samples obtained from wild-type (WT) and αGal knockout (KO) pigs after the explant from the mice subcutaneous area [19,20]. The αGal KO pig is a genetically manipulated αGal-deficient animal in which the gene responsible for the synthesis of the enzyme α1,3-galactosyltransferase (GGTA1, which catalyzes the αGal saccharide and proteins/lipids bond formation) has been silenced. This genetic modification generates a sort of “humanized” animal that is no longer able to synthesize the αGal similarly to humans thus acquiring in turn the ability to produce anti-Gal antibodies. However, this previous approach [19,20] is inherently limited as the implantation of biomaterials in WT animals (constitutively expressing the αGal), is precluding any immunological reaction towards the αGal antigen itself [21]. Considering what has been reported so far, it seems rational to use genetically manipulated αGal-deficient animals, such as GGTA1-KO mice, as recipient animal models. This could mimic the human immunologic environment and, to our knowledge, it has not been used to test the efficacy of the treatments of commercial BHVs to prevent the αGal-immune mediated calcification seen in clinical practice [10].

The main objective of this study was to evaluate the amount of calcium deposition in isolated leaflets from the commercial Trifecta-GT BHV model (Abbot/St.Jude, Santa Clara, CA, USA) implanted for 2 months in GGTA1 KO mice and compared with a parallel investigation carried out for up to 4 months in WT [22]. Alike and at variance to this parallel report the study was further extended to evaluate the anti-calcification efficacy of a polyphenol-based treatment with specific attention to its ability to mask resident antigens to circulating anti-αGal antibodies. Moreover, we have investigated the level of possible residual αGal-epitope in several tissue districts of the KO-mice considering that a residual amount of αGal epitope reactivity has been recognized in biallelic GGTA1-knockout pig cells and implicated as a possible contributor to chronic rejection of GGTA1^−/−^ organs [23,24].

## MATERIAL & METHODS

All animal experiments and surgical procedures were performed in compliance with the Guide for the Care and Use of Laboratory Animals as published by the US National Institutes of Health (NIH Publication 85-23, revised 1996). The use of a mouse animal model for experimental purposes was authorized by the Italian Ministry of Health: project registration number 17E9C.154; authorization number 542/2020-PR. The GGTA1 KO mouse animal model was cloned and is owned by Biocompatibility Innovation Srl.

### Cloning of C57Bl/6 αGal knockout mice

GGTA1 in mice has many expressed transcripts involving different exons, and in some cases, polyadenylating at different sites (see Figure 1). The goal was to inactivate the gene in a manner that none of these transcripts are made into a functional peptide.

**Figure 1.**
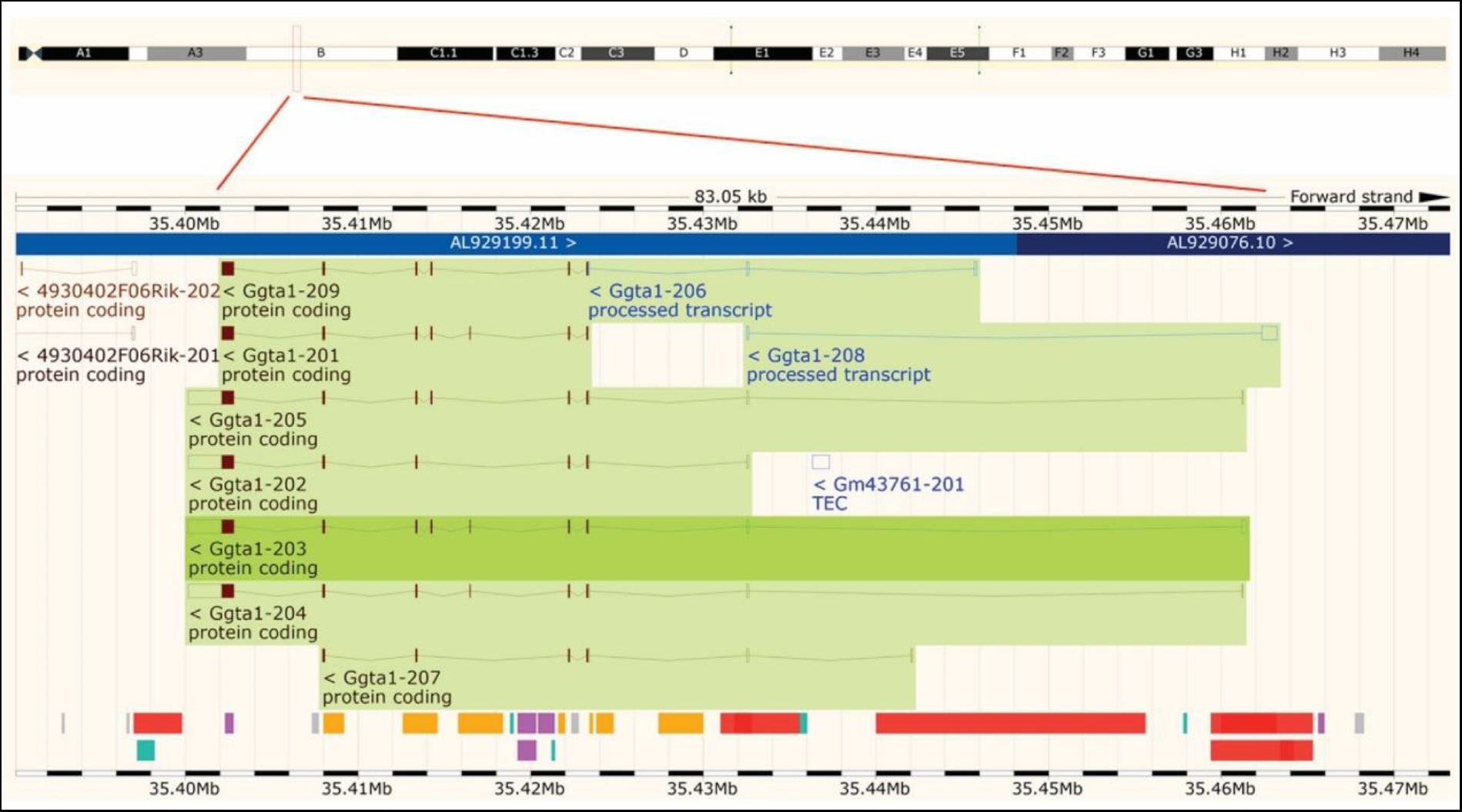
Location of the GGTA1 gene on mouse chromosome 2. The gene is in counter orientation and is represented by several peptide coding transcripts.

The CRISPR/Cas9 system consists of three major elements: a gRNA responsible for site recognition in the genome, the Cas9 protein which introduces the double-strand break in the DNA, and a so-called tracrRNA which mediates the interaction between the gRNA and Cas9 [25]. Together the gRNA and the tracrRNA build up the guide RNA (sgRNA). Pronuclear microinjections were performed using the constructed sgRNA (Figure 2) cloned into an expression plasmid expressing all elements of the CRISPR/Cas9 complex; guide plasmids for both guides were cloned and used.

**Figure 2.**
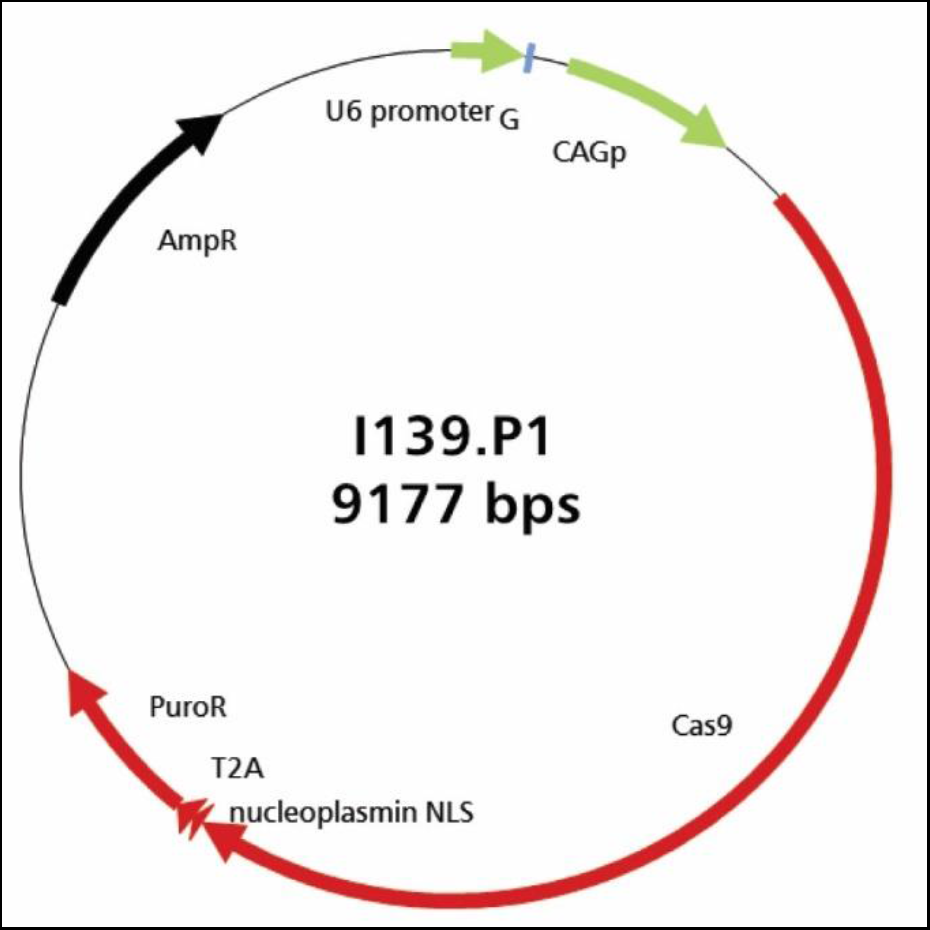
All elements of the CRISPR/Cas9 complex are expressed from this vector: I139.P1 contains the I139.sgRNA1.

Embryonic cells (ES) (60.000/120.000 on 6cm plates) were lipofected (Invitrogen Lipofectamine LTX) using 1μg of the guide plasmid, and 1μg of the homology oligo in each reaction. After 24hs, the cells were selected for 48hs with 0.8μg/ml puromycin (to select for the transient expression of the guide/Cas9 plasmids), and then picked after 8 days (2 x 48 clones from each guide). Candidate clones with visibly reduced amplicon size upon PCR screening were expanded to 24-well format and frozen at −80°C, retested by PCR, and sequenced. Subsequently, appropriate candidate clones were injected into C57Bl/6Ng blastocysts. The injections were highly effective, yielding 80-100% chimeric males from each litter born in three of the five clones. Chimeric mice obtained from the blastocyst injections were mated to C57Bl/6Ng mice to assess transmission to the germ line. Mice were set up for breeding to generate homozygous F2 mice. Finally, the genotype was further reconfirmed.

### αGal quantification in WT e KO mouse tissues

Fresh tissue samples from the different tissues (Table 1) of WT and KO mice were gently blotted on Whatman filter paper, and their weight was recorded (weight range of about 100mg wet weight). Subsequently, they were incubated with the primary anti-αGal antibody M86 [1:50] (mouse; LSBio, Seattle, WA) for 2hs at 37°C with gentle stirring and finally centrifuged at 14.750g for 30min at 4°C.

**Table 1.** Comparison of the αGal quantification in different tissue districts of wild-type (WT) and GGTA1 KO (KO) mice animal model with the relative percentage of αGal silencing (n=7 for each tissue district).

| n° of alpha-Gal epitope/10mg of tissue |  |  |  |
| --- | --- | --- | --- |
| **Tissue/Organ** | **WT** | **KO** | **% of αGal silencing** |
| Eye | $7.68 \cdot 10^{11} \pm 0.08$ | 0 | 100% |
| Thymus | $5.26 \cdot 10^{11} \pm 0.05$ | 0 | 100% |
| Tail | $4.13 \cdot 10^{11} \pm 0.05$ | 0 | 100% |
| Spleen | $1.70 \cdot 10^{11} \pm 0.02$ | 0 | 100% |
| Myocardium | $1.52 \cdot 10^{11} \pm 0.02$ | $2.08 \cdot 10^{10} \pm 0.01$ | 86.3% |
| Kidney | $1.53 \cdot 10^{11} \pm 0.03$ | $2.76 \cdot 10^{10} \pm 0.02$ | 82% |
| Lung | $2.93 \cdot 10^{11} \pm 0.03$ | $5.86 \cdot 10^{10} \pm 0.02$ | 80% |
| Liver | $2.05 \cdot 10^{11} \pm 0.06$ | $4.1 \cdot 10^{10} \pm 0.03$ | 80% |
| Skin | $1.72 \cdot 10^{11} \pm 0.01$ | $9.55 \cdot 10^{10} \pm 0.05$ | 44.4% |
| Brain | $3.35 \cdot 10^{10} \pm 0.02$ | $1.87 \cdot 10^{10} \pm 0.02$ | 44.2% |

The number of αGal epitopes was quantified through a patented ELISA test [26]. Briefly, a Polysorp 96-well plate (Nunc, Rochester, NY, USA) was coated with 100μl of alpha-Gal/HSA (human serum albumin; Dextra Laboratories, Berkshire, UK), 5μg/ml, for 2hs at 37°C. After washing three times with PBS, the blocking procedure was performed using 300μl per well of 2% HSA (Sigma, St. Louis, MO, USA) in PBS for 2hs, at room temperature in darkness. Wells were then washed three times as mentioned above. A set of wells was loaded with 100μl of supernatant derived from tissue samples of wild-type (WT) and KO mice and incubated overnight at 4°C in darkness. After washing, the secondary HRP-conjugate antibody [1:500] (Dako Cytomation, Glostrup, Denmark) was loaded. Finally, 100μl of horseradish peroxidase substrate buffer was added to each well for 5min at room temperature in darkness. The plate absorbance was measured by a plate reader at 450nm (Multiscan Sky, Thermo Scientific). The number of epitopes was calculated by comparison with a calibration line obtained using rabbit red blood cells [27].

### Polyphenols-based treatment of commercial pericardial leaflets

Briefly, a blend of polyphenols was solubilized in phosphate buffer solution (PBS, 50mM NaH_2_PO_4_, 20mM Na_2_HPO_4_) at room temperature as previously described [28–31]. Bovine pericardial leaflets isolated from the commercial Trifecta-GT BHV model (Abbott, Plymouth, MN, USA) were allowed to briefly drain, rinsed with PBS and transferred to the polyphenolic reagent solution, and left to react under moderate constant stirring for two consecutive steps of 30min each, at room temperature in the dark. At the end of incubation, the samples were subjected to two washes in isotonic phosphate buffer for 15min each and stored at 4°C in PBS until the moment of implantation. Samples subjected to the polyphenols-based are labeled as F.

### Mice subcutaneous implantation

Alike and at variance to the parallel investigation [23], GGTA1 KO mice (C57BL/6, 6 weeks old, 30g) instead of WT mice were used. After anesthetizing and shaving, a subcutaneous pouch was created in the dorsal area for each mouse. Each not-treated (NT, n=16) and polyphenols-treated (F, n=16) Trifecta-GT leaflet was implanted into the pouch of each animal, and the wounds were closed with 6/0 nylon sutures. After 1, and 2 months the mice were sacrificed under a CO_2_ atmosphere and the samples were carefully harvested.

### Evaluation of anti-αGal antibodies production in KO mice

Anti-αGal serum IgM and IgG antibodies from the αGal KO mice were determined before and 2 months after implantation of original and polyphneols-treated leaflets form Trifecta-GT, by enzyme-linked immunosorbent assay (ELISA). About 0.5 – 1.0ml of blood per mouse was collected by infraorbital venous plexus sampling (n=10). A Polysorp 96-well plate (Nunc, Rochester, NY, USA) was coated with 100μl of alpha-Gal/HSA (Bovine serum albumin; Dextra Laboratories, Berkshire, UK), 5μg/ml, for 2hs at 37°C. After washing three times with PBS, the blocking procedure was performed using 300μl per well of 2% HSA (Sigma, St. Louis, MO, USA) in PBS for 2hs, at room temperature in darkness. Wells were then washed three times as mentioned above. A set of wells was loaded with 100μl of [1:80] diluted serum and incubated overnight at 4°C in darkness. After washing, the secondary HRP-conjugate anti-mouse IgM and IgG antibody [1:500] (Jackson Immunoresearch, Pennsylvania, USA) were loaded. Finally, 100μl of HRP substrate buffer was added to each well for 5min at room temperature in darkness. The plate absorbance was measured by a plate reader at 450nm (Multiscan Sky, Thermo Scientific).

### Calcium quantification in explanted commercial leaflets

Polyphenols-treated (F) and non-treated (NT) leaflets from Trifecta-GT BHVs were carefully explanted from KO mice and washed twice in sterile cold PBS for 10min. Specimens were subsequently subjected to acid hydrolysis in HCl 6N at 110°C for 12hs. Calcium evaluation was performed in hydrolyzed samples by inductively coupled plasma according to the directives of the EPA6010D method [32] and expressed as μg Ca^2+^/10mg of dry defatted weight (ddw). As a control sample, calcium quantification was also carried out in unimplanted off-the-shelves original Trifecta GT™ valve leaflets.

Ddw was determined by comparing lyophilized dry-weight samples before and after delipidation treatment. After the lyophilization step, sample tissues were incubated for 36hs under 10 kPa over P_2_O_5_ at 37°C until a constant dry weight was attained. The defatted procedure was carried out by incubation of tissue specimens in ascending series of alcohols followed by two steps of chloroform/methanol (2:1 and 3:1), in a descending series of alcohols, and finally in the water.

### Von Kossa staining in explanted commercial leaflets

Representative polyphenols-treated (F) and non-treated (NT) tissue samples explanted from WT (at 4 months of follow-up, n=4) and KO (after 2 months of follow-up, n=4) mice were carefully rinsed with cold PBS and subsequently embedded in OCT compound (Tissue Tek; Sakura Finetek, Tokyo, Japan), cryo-cooled in liquid nitrogen, and cut into 8μm cryosections. Sections were stained with Von Kossa. The general appearance of the extracellular matrix (ECM) and calcium deposition were examined.

### Statistical analysis

The data were analyzed in Microsoft Excel® and Prism® 7 for Windows (v7.03, GraphPad Software lnc., California) and expressed as mean ± standard deviation (SD). A two-sided unpaired T-test was used to assess significant differences between the treated and untreated groups, at the 0.95 confidence level.

## RESULTS

### αGal quantification in WT e KO mouse tissues

As reported in Table 1, the GGTA1 gene silencing inhibited αGal antigen synthesis in a non-uniform manner, ranging from 100% to 80%, depending on the tissue district. For both the skin and the brain samples, the inhibition of the antigen expression was limited to 44%. Interestingly, the brain of the WT mouse exhibits the lowest number of αGal antigens while accounting for the same epitope amount determined in the KO mouse after gene silencing.

### Evaluation of anti-αGal antibodies production in KO mice

As a result of food intake during housing, the GGTA1 KO mice develop a bacterial flora expressing the αGal antigen, thus leading to the onset of a baseline level of IgG and IgM anti-αGal antibodies. This baseline level appeared to be further increased following the implantation of the Trifecta-GT leaflets (comprising glutaraldehyde-treated bovine pericardium) previously reported to contain a significant amount of this antigen [11]. In this specific case, the 2 months of permanence of the implants in the mouse subcutis, doubled the level of circulating anti-Gal IgG and more than tripled that of IgM (Figure 3, BL vs NT). Particularly, the polyphenol-based treatment demonstrated the ability to make the treated tissue “undetectable” to the mouse’s immune system while preventing the increase of anti-Gal antibody levels otherwise observed in the case of NT samples implantation: there were no statistically significant differences between the IgG and IgM baseline levels found after 2 months of implantation of the polyphenols-treated leaflets (Figure 3, BL vs F).

**Figure 3.**
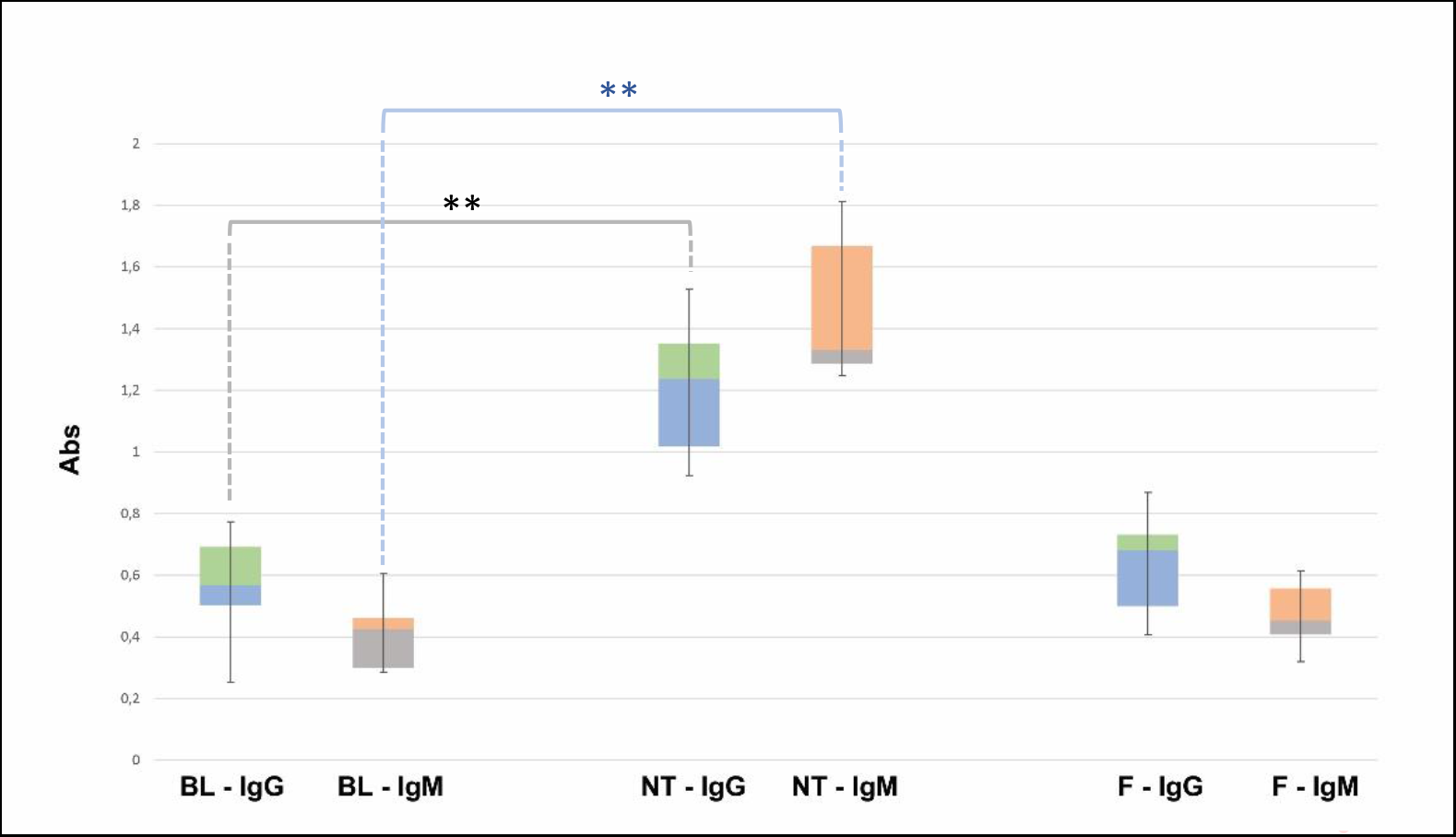
Quantitative evaluation (450 nm OD absorbance units) of IgG and IgM anti-Gal antibody production in KO mice. On the left the basal level (BL), in the center, and on the right the variations found after 2 months of implantation of commercial Trifecta-GT valve leaflets not-treated (NT) and polyphenols-treated (F) (n=10 for each type of sample including BL). The data points represent the means ± SD. ** Indicates a statistically significant difference between the two groups at the 0.95 confidence level.

### Calcium quantification in explanted commercial BHV leaflets

In Trifecta-GT leaflets explanted from the KO mice, a relevant calcium deposition (Figure 4, grey bar) was evident even after one month and accounted for more than four times the amount found in the WT mouse at the same time (1 month KO *vs* 1 month WT p=0.015). The intensity of mineralization increased after two months, it was not significantly different from that determined in the WT mouse at the same time and comparable to that determined in the 4-month WT model. Again, similarly to what was already evidenced in the WT mouse, the polyphenols-based treatment exhibited a strikingly evident anti-calcification effect even in the KO-F samples. The surprisingly efficient treatment with polyphenols appears to prevent by over 99% the calcium deposition in both WT and KO animal models.

**Figure 4.**
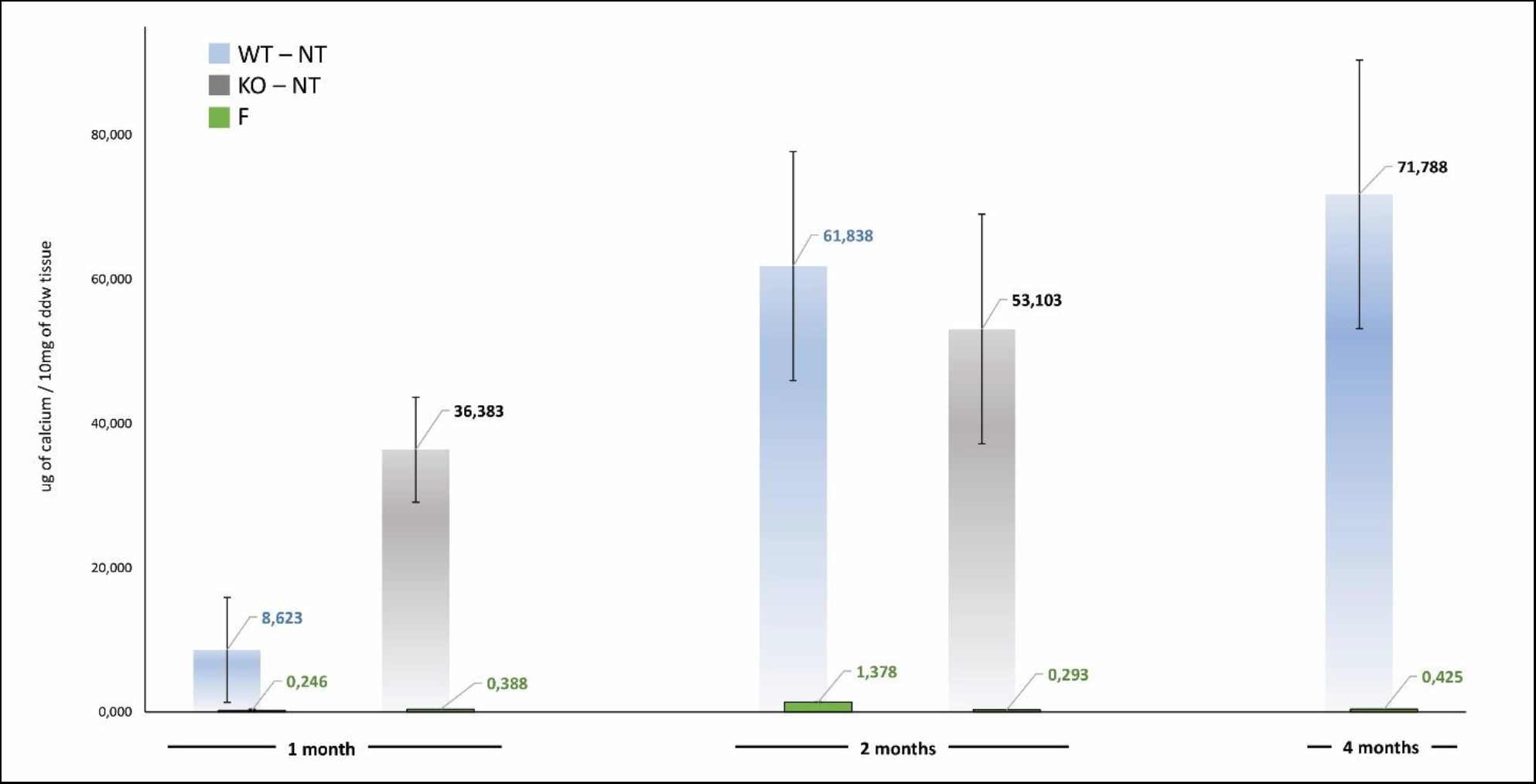
Calcification trend in not-treated (NT) and polyphenols-treated (F, green bar) currently adopted leaflets of Trifecta-GT implanted in the subcutis back area of wild-type mice (WT, light blue bar at 1, 2, and 4 months of follow-up) and knockout for αGal antigen (KO, grey bar at 1 and 2 months of follow-up). As a control sample, calcium quantification was also carried out in un-implanted off-the-shelves original Trifecta GT™ valve leaflets resulted to be 1.19 ± 0.05 μg/10mg of ddw.

### Von Kossa staining in explanted commercial leaflets

The histological evaluation of calcium deposition, in some representative explanted leaflets (Figure 5), was found to be in line with the results of plasma analysis (Figure 4). In general, the non-treated leaflets (NT) exhibited unevenly diffused micro-calcifications. Their counterparts, treated with polyphenols (F), did not show calcified spots even when calcium content accounted for about 0.3 μg/10mg of ddw. It is known that the sensitivity of the quantitative Inductive Coupled Plasma technique is considerably higher than the histological evaluation. Particularly, the calcium content of the F leaflets resulted below the detection limit of the Von Kossa staining, besides the fact that, unexpectedly, it was even significantly lower than that determined in the untreated control group samples before implantation.

**Figure 5.**
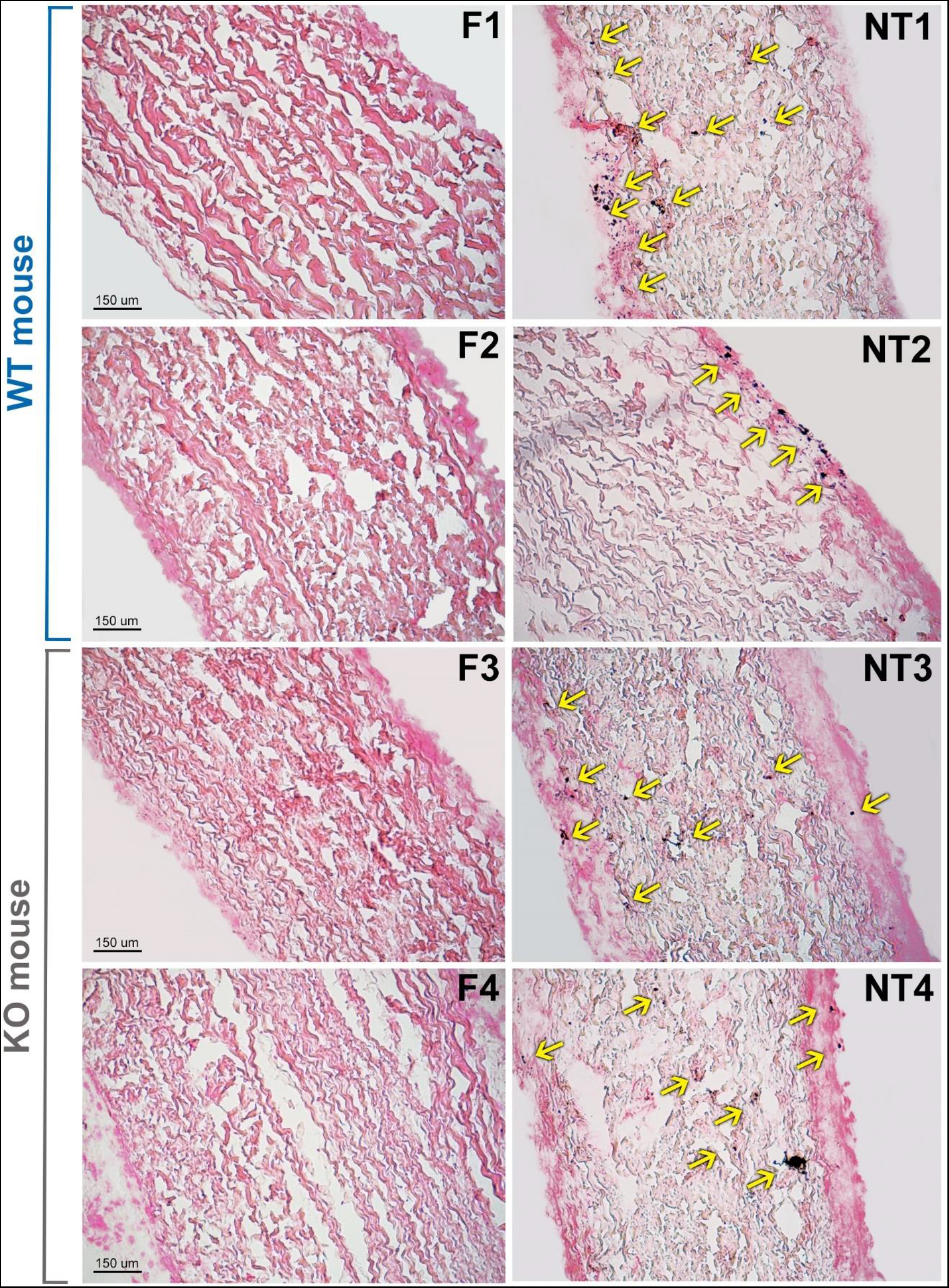
Histological evaluation of calcium deposition in representative not-treated (NT) and polyphenols-treated (F) leaflets from Trifecta-GT valve implanted in the subcutis back area of wild-type (WT, 4 months of follow-up) and αGal knockout (KO, 2 months of follow-up) mice. Spots of calcified deposition are highlighted by yellow arrows. Von Kossa staining, magnification 10X.

## DISCUSSION

Recently, the results of the Translink international collaborative study group have been released [10]. Translink is a prospective European Union-funded collaborative project, which assessed the role of the xenoantigens in BHVs deterioration. In particular, Translink is focused on the involvement of anti-glycan antibodies in inducing calcification of BHV tissues, suggesting that BHV xenogeneic antigens contribution to the immunogenicity of animal-derived implants, is eliciting antibodies that are likely involved and support valve calcification. The results obtained from this study provide evidence that the lack of the αGal epitope in the GGTA1 KO mice was associated with an early (one-month) response leading to quadrupling the calcium deposition rate determined in WT animals at the same time (Figure 4). This initial response is recalling the severe calcific deposits associated with the early failure of porcine heart valve transplanted in pediatric patients [33] and successively suggested to be related to the presence of residual αGal epitopes [34].

In agreement with what was reported in humans by Senage T and colleagues [10], (tissues in which masked xenoantigen carbohydrates, do not trigger antibody-mediated calcification), the anti-αGal antibodies in the GGTA1 KO blood analysis resulted in a remarkable IgG and IgM increase only in mice that received a bioprosthetic leaflet not treated with polyphenols (Figure 3). Accordingly, the use of polyphenols results in a powerful approach able to prevent calcium deposition as assessed in two different mouse animal models. Of interest that polyphenols have also been reported to inhibit calcium deposition in BHVs when tested in an in vitro system [28].

Small animal models such as rats or mice are widely used for in vivo biomaterial assessment for their low cost, ready availability, ease of handling, and well-defined immune parameters. These models are generally used for the assessment of chronic changes to BHV leaflets implanted in an ectopic (non-cardiac) location. In particular, the subdermal model provides permanent contact of the implant to host tissue and sufficient blood supply (serum exposure), which eases cellular infiltration and allows a rapid screening efficacy for anti-calcification treatments.

Particularly, the use of the GGTA1 KO animal model enables a differential evaluation of the immune-mediated effects of the αGal concerning that of the whole of other intrinsic and extrinsic factors leading to the calcification of BHVs (as also revealed by the WT mouse model). In fact, besides the early calcium deposition, the increase of anti-αGal IgG and IgM, specifically due to the presence of the αGal epitope in the implanted tissues, is opening the way to the separate detection of residual αGal antigen in any kind of implantable biomaterials. In addition, the results of this investigation are further confirming the presence of immunologically active αGal antigens in BHVs currently adopted in clinical practice as previously determined, by a different analytical approach [11].

Formation of the αGal epitope mostly occurs by the transfer of galactose in a α(1,3)-glycosidic linkage to an N-acetyllactosamine (LacNAc) acceptor molecule group presented on protein and lipid (Figure 6). This reaction is encoded by the GGTA1 gene [35]. However, a residual amount of αGAL epitope reactivity has been recognized in biallelic GGTA1-knockout pig cells and implicated as a possible contributor to chronic rejection of GGTA1^−/−^ organs as found in non-primate models of xenotransplantation [23,24]. The existence of another Gal-transferase in GGTA1 KO mice was already reported by Milland et al. [36] identifying isoglobotrihexosylceramide synthase (iGb3s) in GGTA1 KO mice and the presence of a small amount of iGb3s in tissues using monoclonal antibodies. IGb3S is known to generate the Galα1,3Gal disaccharides epitopes on glycosphingolipids (Lac) by the addition of galactose to lactosylceramide (Figure 6).

**Figure 6.**
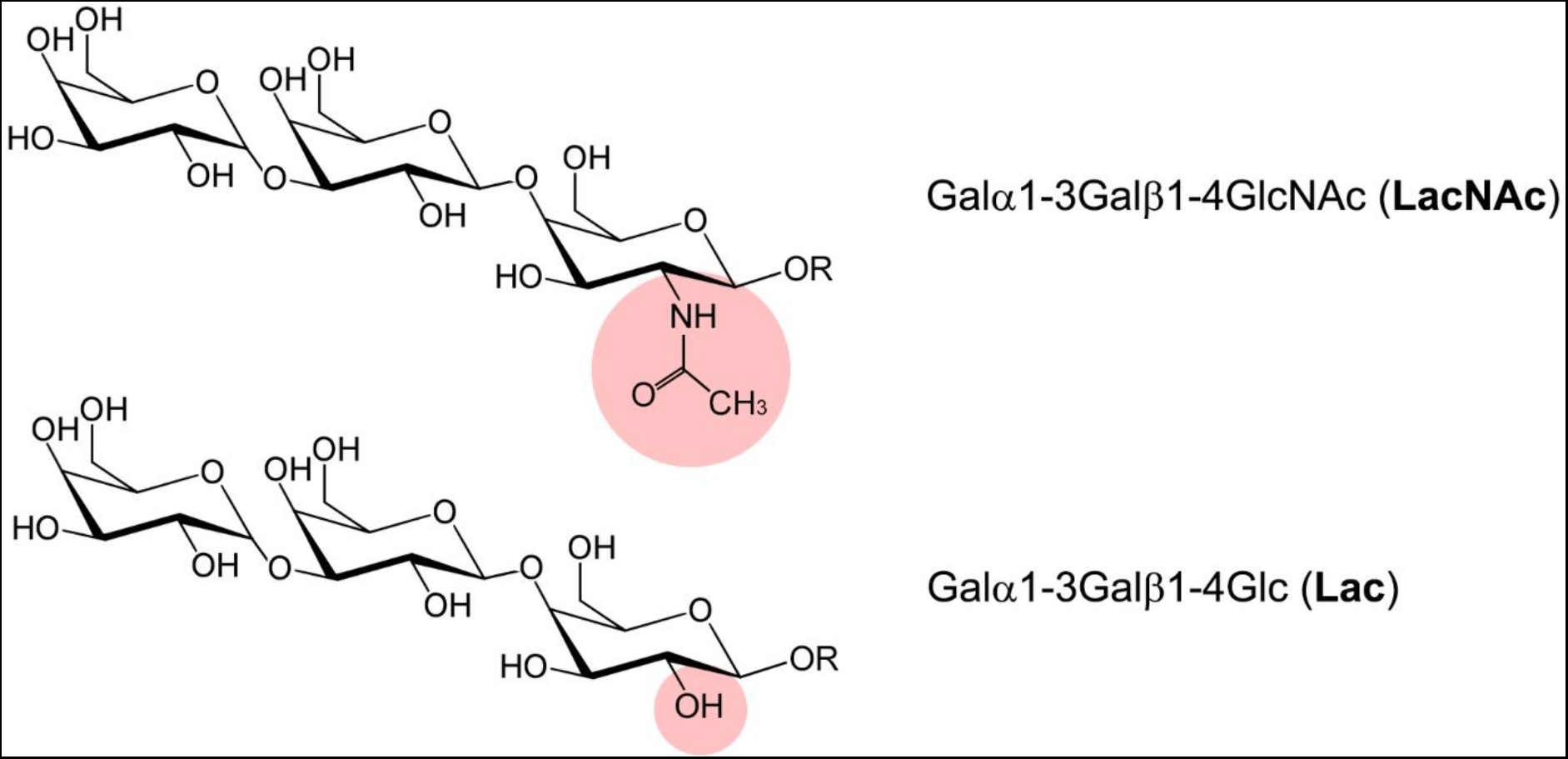
Structure of the different αGal xenoantigenic trisaccharides, at the top NAc and the bottom Lac. The bonds responsible for the different origins of the antigen are highlighted in red.

The comparison of the αGal epitopes number as quantified in the different tissues of WT and GGTA1 KO mice confirmed what has already been reported in the literature namely the presence of variable antigenic residues despite the silencing of the GGTA1 enzyme (Table 1). To our knowledge, this is the first report, to provide the distribution of the residual percentage of αGal likely related to the enzyme iGb3s compared to the uppermost activity of GGTA1.

The αGal epitope is related to the iGb3s enzyme exclusively bound to lipid components, in particular, to ceramides. In the mammalian nervous system, nerve conduction is facilitated by myelin, a lipid-rich membrane that wraps around the axon. The myelin sheath is a specialized structure with distinct lipid and protein constituents. Galactosylceramide (GalCer) and sulfatide make up approximately 30% of total myelin lipids [37]. In particular, the levels of GalCers are especially high in the brain and have been reported to be higher than glucosylceramides (GluCers) in the WT mouse brain [38]. Previous studies revealed that ceramides mediate signal transduction and cell adhesion and are crucial for the formation of nervous tissues [39], this could explain why even in the KO mouse a significant amount of iGb3s-αGal epitopes are available, due to their unavoidable presence for correct brain and neuronal function. The outermost layer of the mammalian epidermis is the stratum corneum, which is made of flattened, enucleated keratinocytes and a unique extracellular lipid matrix produced by differentiating keratinocytes. The stratum corneum provides the permeability barrier against water and various environmental agents, such as chemicals and microorganisms. About half of the lipids in the stratum corneum are mixtures of ceramides [40]. Significant levels of ceramides were also found in the kidneys, liver, lungs, and myocardium [41]. However, the residual percentages of αGal, quantified in this study, agree with what was reported by Shao A and colleagues who describe a reduction in the expression of the antigen between 5.19% and 21.74% in GGTA1-KO mice [21], except for the skin and brain areas where, due to a higher concentration of ceramides, the reduction is much smaller.

Noteworthy, GGTA1 KO animals do not seem to express iGb3s in sufficient amounts to mediate cell destruction: the work of Murray and colleagues reports as a minimum threshold of αGal expression is required to induce antibody-mediated skin graft rejection in a mouse GGTA1 KO model [43]. Accordingly, a previous study demonstrated that silencing the porcine iGb3S gene did not affect measures of anticipated pig-to-human and pig-to-primate acute rejection, suggesting iGb3S is not a contributor to antibody-mediated rejection in pig-to-primate or pig-to-human xenotransplantation [43]. In fact, the αGal contribution due to the presence of iGb3S is not appreciable by heat-inactivated human and baboo sera antibodies when incubated with GGTA1 KO or GGTA1/iGb3S double KO pig tissue. This is the reason, even if residues of αGal antigen are still present, the αGal KO mouse was revealed as an adequate animal model for evaluating the calcification propensity of currently adopted BHV tissues. The implantation of Trifecta-GT valve leaflets has significantly stimulated the mouse immune system and resulted in massive production of anti-Gal antibodies (Figure 3) both IgM and IgG type. This mouse immune-mediated reaction is apparently quite similar to that occurring in humans, as extensively reported in the literature [12,13].

All that raises important considerations in evaluating the efficacy of anti-calcific treatments, making it clear that the choice of small animal models must necessarily prefer the KO model in order not to incur a possibly dramatic underestimation of the potential triggered by immune-mediated reactions towards xenoantigens. What is more, the polyphenol-based treatment applied in this investigation, in addition to having recently been proven effective in inhibiting the extrinsic mechanisms related to commercial BHVs degeneration [22], showed an unexpected ability to inhibit the recognition of BHV xenoantigens by circulating antibodies in a GGTA1 KO mouse while completely preventing calcific depositions compared to the untreated counterpart.

## DISCLOSURES

FN, AG, and GS work for Biocompatibility Innovation Srl holding the position of Chief Executive Officer, Chief Technology and Innovation Officer, and Laboratory Manager respectively.

## Notes

### Competing Interest Statement

The authors have declared no competing interest.

